# Epistasis and cryptic QTL identified using modified bulk segregant analysis of copper resistance in budding yeast

**DOI:** 10.1101/2024.10.28.620582

**Authors:** Cassandra Buzby, Yevgeniy Plavskin, Federica M.O. Sartori, Qiange Tong, Janessa K. Vail, Mark L. Siegal

## Abstract

The contributions of genetic interactions to natural trait variation are challenging to estimate experimentally, as current approaches for detecting epistasis are often underpowered. Powerful mapping approaches such as bulk segregant analysis, wherein individuals with extreme phenotypes are pooled for genotyping, obscure epistasis by averaging over genotype combinations. To accurately characterize and quantify epistasis underlying natural trait variation, we have engineered strains of the budding yeast *Saccharomyces cerevisiae* to enable crosses where one parent’s chromosome is fixed while the rest of the chromosomes segregate. These crosses allow us to use bulk segregant analysis to identify quantitative trait loci (QTL) whose effects depend on alleles on the fixed parental chromosome, indicating a genetic interaction with that chromosome. Our method, which we term epic-QTL (for *<u>epi</u>*static-with-*<u>c</u>*hromosome *<u>QTL</u>*) analysis, can thus identify interaction loci with high statistical power. Here we perform epic-QTL analysis of copper resistance with chromosome I or VIII fixed in a cross between divergent naturally derived strains. We find seven loci that interact significantly with chromosome VIII and none that interact with chromosome I, the smallest of the 16 budding yeast chromosomes. Each of the seven interactions alters the magnitude, rather than the direction, of an additive QTL effect. We also show that fixation of one source of variation — in this case chromosome VIII, which contains the large-effect QTL mapping to *CUP1* — increases power to detect the contributions of other loci to trait differences.

**Author Summary:** Most traits of interest that vary in populations are determined by multiple genetic factors, as well as by environmental variation and random chance. These influences may combine in complicated ways, for example when the effect of one genetic variant depends on genetic variants elsewhere in the genome. Such dependencies are difficult to identify and characterize because testing many possible combinations of influences reduces statistical power. We address this challenge by combining bulk segregant analysis, a robust method for detecting effects of individual genetic variants averaged across genetic backgrounds, with chromosome fixation in budding yeast. This approach allows us to detect gene-variant effects that depend on the fixed chromosome with statistical power comparable to that achieved when detecting background-independent effects. Applying the approach to copper sulfate resistance, we identify and characterize interaction effects and, by removing one source of variation (a single yeast chromosome), improve detection of background-independent effects as well.

## Introduction

Predicting phenotype from genotype is a central goal of genetics research, yet even with great advances in sequencing technology and experimental scale, accurate characterization of how genetic differences translate into trait differences remains elusive. Many genetic influences are subtle, and most traits of interest that vary in populations are determined by multiple genetic factors, as well as by environmental variation and random chance [1–4]. Moreover, these influences may combine in complicated ways, for example in the form of gene-by-gene (epistatic) or gene-by-environment interactions [5–7].

Epistasis is particularly challenging to detect in standard quantitative-genetic analyses because the number of possible interaction effects vastly exceeds the number of single-locus (additive) effects, leading to very low statistical power to detect the interactions [5]. Standard analyses fit additive effects before interaction effects, which can lead to the mistaken inference that epistasis is less salient than it actually is [8,9]. If interacting alleles are rare in a mapping population, under-detection of epistasis is even more severe. Failure to detect epistasis in turn compromises accurate prediction of an individual’s phenotype from genotype, even when an additive model appears to fit well [8–10].

The distribution of types of epistatic effects influences both detection of loci with effects on traits and evolutionary trajectories [11–13]. Magnitude epistasis changes the size of an allele’s effect without changing its direction (beneficial vs. detrimental), whereas sign epistasis changes the direction. Sign epistasis gives rise to peaks and valleys in fitness landscapes and can alter the order of mutations that increase fitness [11]. Models of distribution and patterns of these effects have been postulated, with two extremes being global epistasis and idiosyncratic epistasis. Global epistasis indicates underlying predictability of the effects of individual alleles, perhaps based on a latent phenotype such as protein stability or physiological health [14,15]. Idiosyncratic epistasis, by contrast, refers to unpredictable allele effects that ultimately depend on details of particular alleles [16,17]. Data on the relative prevalences of magnitude and sign epistasis, which are currently lacking, would inform which model of epistasis patterns is more suitable, which in turn will inform models of the evolution of complex traits.

An ideal method for identifying and characterizing epistatic contributions to trait variation would have equal (and high) statistical power to detect additive and interaction effects. Experimental designs with extremely large sample sizes are now common [18–22], but the most powerful of these favor (or exclusively detect) additive effects. One such experimental design is bulk segregant analysis (BSA). In a typical BSA experiment, many progeny are produced from a cross between divergent parental genotypes, progeny with extreme values of a trait of interest are pooled, and high-coverage genome sequencing is used to detect loci at which the allele frequency in the selected bulk deviates significantly from the allele frequency in an unselected (or oppositely selected) bulk [18,23]. Although BSA benefits from sampling many individuals — over 10 million in some cases [18] — it cannot detect genetic interaction effects because pooled sequencing results in an estimate of the effect of any one locus that is the average across all genotypes at other loci.

Genotyping and phenotyping individuals rather than pools avoids the averaging issue, but suffers from the problem that potential pairwise interactions far outnumber additive effects of single loci, resulting in greatly reduced statistical power. Two recent attempts to detect epistasis using very large sample sizes illustrate this challenge. One study, which genotyped ∼200,000 diploid progeny of a yeast cross, identified numerous interaction effects, but found a strong positive correlation between the additive effect of a locus and the number of pairwise interactions it participates in [22]. This correlation could reflect the true distribution of epistatic effects or an influence of additive-effect size on detectability of interactions [22]. Another study genotyped ∼100,000 haploid segregants but to maximize power tested only interactions between pairs of additive-effect QTL [21]. This study found evidence of epistatic effects, but found the additive-plus-pairwise model to have only modest predictive power for the phenotypes of reconstructed genotypes, suggesting that effects remain undetected [21]. Another recent study did not genotype large numbers of segregants, but instead engineered naturally occurring gene variants into four divergent yeast strains to detect genetic background-dependent effects on growth. This study found a substantial fraction of variants to have background-dependent effects, but was limited to variants in and around 103 genes previously shown to be interaction hubs in genome-wide double-mutant screens [24,25]. The results of these studies suggest that unbiased approaches are needed that equalize the power to detect additive and epistatic effects.

To identify and characterize genetic interactions with comparable statistical power to that of additive effects, we combine chromosome substitution with BSA in the genetically tractable budding yeast, *Saccharomyces cerevisiae*. We call this method epic-QTL (for *<u>epi</u>*static-with-*<u>c</u>*hromosome *<u>QTL</u>*) analysis. To perform epic-QTL, we cross two divergent natural isolates of *S. cerevisiae*, but before generating segregating F2 progeny from the resulting F1 diploid, we first eliminate one chromosome from one parent, fixing the remaining homologous chromosome. The result is that all surviving F2 haploids contain identical copies of one chromosome, but have unique combinations of the two parents’ alleles on all other chromosomes. We also perform the reciprocal cross, in which the other parent’s chromosome is fixed. We then grow the haploid progeny in selective or non-selective conditions. Here, we use the presence or absence of copper (II) sulfate (CuSO_4_) as a selective condition, with the presence of copper and subsequent oxidative stress selecting for individuals with alleles that promote survival in this environment. Standard BSA would compare the allele frequencies in bulks grown in the presence or absence of copper to identify additive QTL for the trait of copper resistance. In epic-QTL, there are four bulks rather than two. By comparing the QTL identified in the crosses with the reciprocal fixed chromosomes, we can identify loci whose effects differ between genetic backgrounds. The loci that are dependent on parental-chromosome background for their effect are thus epistatic with that fixed chromosome and are detected with statistical power usually only afforded to additive effects. In practice, this comparison is done by including a parental-chromosome (or genetic-background) term in a logistic-regression model that is fit to the data from all four bulks simultaneously. An additional benefit of chromosome substitution is an increase in power for detection of additive effects by removing a source of variance in a population. With only one cross, the fixation of a chromosome containing a major QTL allows the remaining segregating sites to exert greater influence over membership in the selected bulk.

Using epic-QTL we can determine where QTL that interact with each fixed chromosome lie relative to where additive QTL lie, and how prevalent interactions are for a given trait, in this case copper sulfate resistance. We focus on copper sulfate for three reasons: 1) its extensive use in prior *S. cerevisiae* studies, allowing comparison with previous results; 2) the potential for unexplained heritability, in that copper resistance shows a large gap in broad and narrow sense heritability compared to other traits [4]; and 3) its ecological and physiological relevance [26–28]. In addition, one major interaction hub identified by Matsui *et al.* was *CUP1* on chromosome VIII, which is a major driver of copper resistance, and thus a perfect candidate for chromosome fixation [22]. We fixed chromosomes I or VIII in a cross between strains derived from an oak tree or from a wine barrel, which differ on average at 1 in 200 base pairs [29]. The variants in these strains reflect natural genetic diversity, offering insights into genetic interactions that might occur within populations. We demonstrate that this method can be used to remove sources of trait variation to reveal additive QTL otherwise undetected at this experimental scale, as well as to identify interaction QTL with high statistical power. We find that these interactions often occur within significant additive QTL peaks, providing evidence of a lack of pervasive sign epistasis.

## Results

### Implementation of epic-QTL for copper resistance

Our method to identify QTL interacting with a fixed chromosome, or epic-QTL, combines chromosome substitution with BSA in a cross of two parental yeast strains. For the parental strains we used divergent natural isolates of *S. cerevisiae*: NCYC3631 and NCYC3591 (hereafter “Oak” and “Wine”, respectively). Before crossing the two strains, we engineered one strain of each parent to contain the inducible *GAL1* promoter (*pGAL1*) adjacent to the centromere of one chromosome. Inducing a high level of transcription through the centromere destabilizes the kinetochore, resulting in missegregation of the chromosome during mitosis so that a strain with only the homologous chromosome remaining can be isolated [30]. By this method, we fixed chromosome I in each background (Fig 1A) and, separately, chromosome VIII in each background (Fig 1B). We then mated the Wine *pGAL1* strain with the Oak non-*pGAL1* strain (Fig 1C), and vice versa, to produce diploids with reciprocal conditional chromosomes. Each diploid strain was then plated on galactose to induce *pGAL1*. After re-plating colonies from the galactose-containing media to ensure clonality, individual colonies were checked by Sanger sequencing at biallelic regions to confirm loss of the conditional chromosome. Each of these aneuploid (2N–1) strains was then sporulated to produce an F2 population of haploid segregants fixed for a single parent’s chromosome (Fig 1C), which could then be used for BSA (Fig 1D).

**Fig 1.**
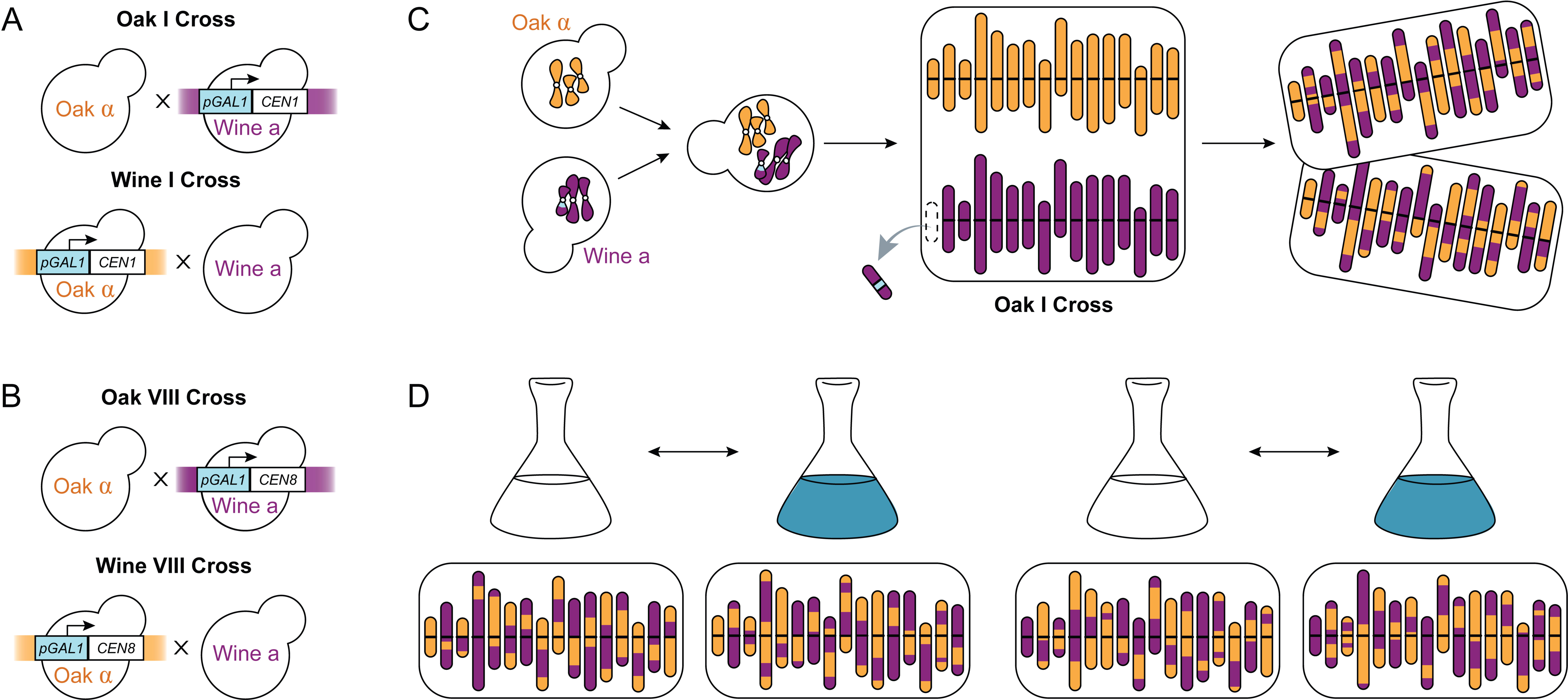
Insertion of *GAL1* promoter allows for fixed chromosomes. (A–B) Matings to produce reciprocal diploids with single conditional chromosomes for chromosome I (A) and chromosome VIII (B). (C) Induction of the *GAL1* promoter adjacent to the centromere destabilizes the chromosome’s kinetochore, leading to loss of that chromosome in diploids and fixation of the homologous chromosome in F2 haploid progeny. One representative cross of the four (Oak I) is shown. (D) BSA experiment for interactions with chromosome I, allowing detection of additive effects by comparison of selected (blue) and unselected (white) bulks and of interactions with chromosome I background (orange or purple). One representative haploid segregant is shown beneath each bulk’s flask.

Before proceeding to BSA, we demonstrated that the Oak and Wine parental strains, which have been shown to differ phenotypically in multiple traits [31,32], differ in the phenotype of copper sulfate resistance. We grew populations of haploids of each parental strain in an array of doses of CuSO_4_ in rich media (YPD), and compared the cell counts of each population to a control of the same seed culture. The Wine haploid strain showed higher resistance to CuSO_4_, surviving at almost double the percentage at which the Oak haploid strain survived in mid-range doses of CuSO_4_ (S1 Fig).

Due to chromosome fixation, the relative intensity of selection varied at the same doses of CuSO_4_ between crosses. Specifically, as shown below, Wine chromosome VIII has a strongly advantageous allele, so the same dose of CuSO_4_ produces more survivors in the Wine VIII cross than in the Oak VIII cross. We therefore tested multiple doses of CuSO_4_ to determine the dose that produced 1–10% survival of the population. This range of selection optimizes BSA by balancing detectable effects and retention of causative genetic variation in the surviving population [33]. Unselected bulks from each cross were seeded in YPD at 10% the cell number at which selected bulks were seeded so that their final cell densities were approximately the same as those in the selected bulks.

### Known drivers of resistance are identified as additive QTL

Previous studies have identified additive QTL for the trait of copper sulfate resistance in a variety of *S. cerevisiae* strains [2,4,18,34]. In addition, multiple studies have identified mechanistic drivers of copper resistance, including certain alleles of the *FRE1* gene and copy number amplification of the *CUP1* locus [27,35,36]. Because there is overlap across strains for identification of some QTL [37], one form of validation of epic-QTL would be for previously known resistance loci to be detected by epic-QTL as additive effects (that is, effects that do not depend on the parental origin of the fixed chromosome).

In epic-QTL, as in standard BSA, allele frequency differences between bulks at a given genomic site indicate a contribution of variation at that site to bulk membership. For the reciprocal chromosome I crosses and, separately, for the reciprocal chromosome VIII crosses, we sequenced the four bulks (selected vs. unselected for each Oak or Wine fixed chromosome) at high depth of coverage (average ∼70X per site per bulk) to estimate allele frequencies. We partitioned the effects of selection, parental-chromosome background (hereafter “background”), and their interaction using a logistic regression model as described in *Materials and Methods*. The selection term’s coefficient reflects the additive effect on CuSO_4_ survival regardless of parental-chromosome background, whereas the interaction term reflects epistasis with the fixed chromosome. To visualize significant effects across the genome, we use the z-score of each coefficient. The z-score is advantageous over the parametric *p*-value to which it corresponds because its sign indicates the direction of effect of the fixed chromosome, and because we use permutations of it to determine a threshold of false discovery rate, detailed in *Materials and Methods*. To compare significance regardless of effect direction, the majority of plots display absolute values of z-scores. Signed z-scores for each coefficient and position are included in supplemental data (S1 Table).

The additive (selection) peaks on chromosomes VIII and XII are very close to the known copper-resistance genes, *CUP1* and *FRE1*, respectively (Fig 2). These genes are well documented to play critical roles in copper detoxification mechanisms in yeast. *CUP1* encodes a metallothionein involved in sequestering copper ions, thereby mitigating copper toxicity within the cell [38]. *FRE1* encodes a ferric reductase enzyme that contributes to iron uptake and detoxification of copper [39,40]. In both fixed-chromosome backgrounds, we identify *FRE1* as a major driver of resistance, with peak maxima <6 kb away from the coding region (Fig 2). The crosses for the two fixed-chromosome backgrounds are independent, so the agreement between the two crosses further validates the method. In the strains fixed for chromosome I, the peak on chromosome VIII is upstream of the *CUP1* coding sequence. *CUP1* has two paralogs, *CUP1-1* and *CUP1-2*, within a larger tandemly duplicated region in the reference genome of the laboratory strain S288C. We found that the region surrounding these paralogs had poor mapping quality, likely due to repetitive sequences and potential *CUP1* copy variation, which is a known mechanism of resistance [38]. These regions were therefore excluded from our variant calling, and so the peak upstream of the coding sequence is as close as can be to this known driver of copper resistance.

**Fig 2.**
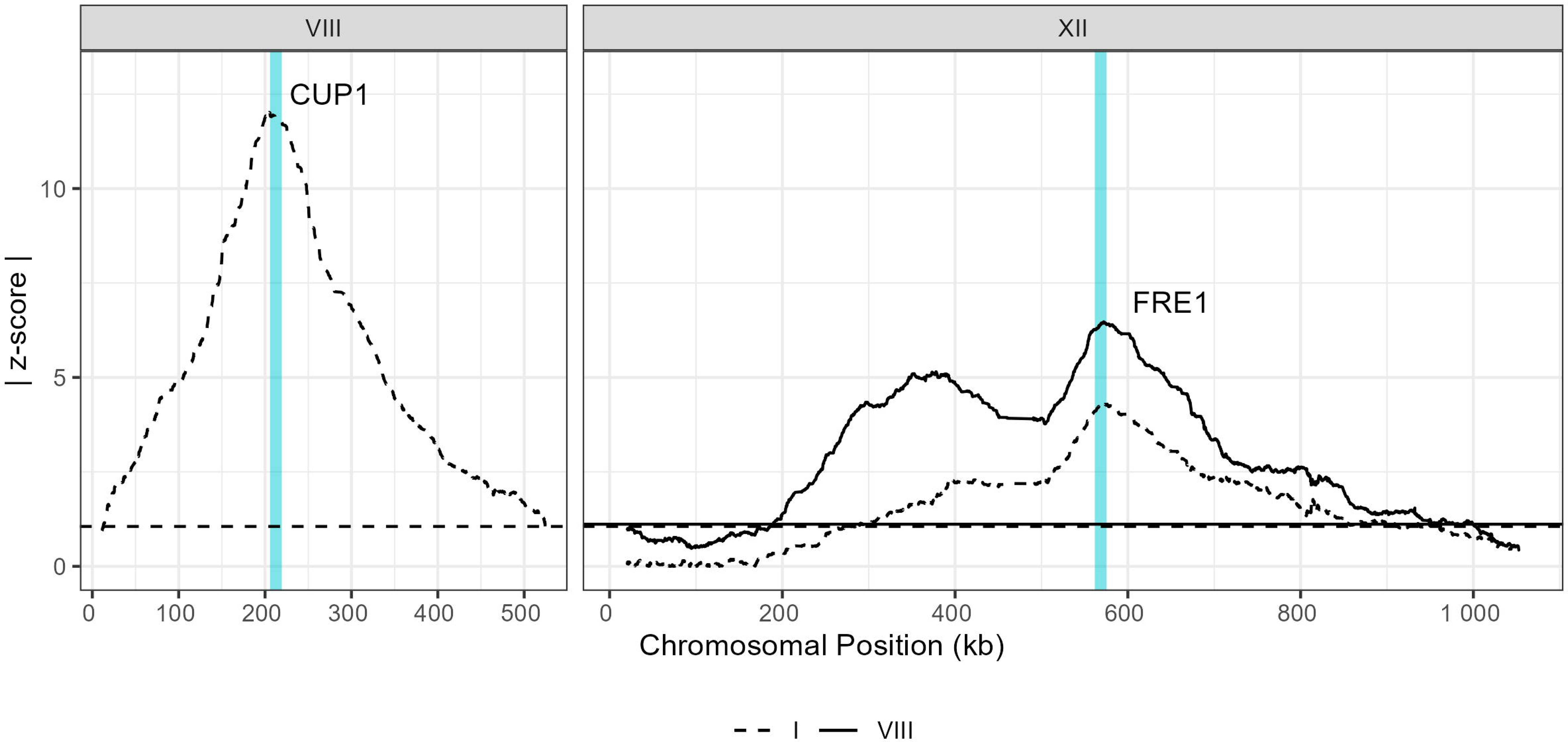
Known drivers of copper resistance identified as additive QTL. Traces represent absolute z-scores of additive (selection) coefficients on chromosomes VIII and XII, as indicated. Dashed line corresponds to the experiment with chromosome I fixed and solid line corresponds to the experiment with chromosome VIII fixed. Horizontal lines indicate the 5% false discovery rate (FDR) threshold for each experiment. Turquoise vertical lines represent the locations of CUP1 and FRE1, both of which fall under large-effect additive peaks as expected.

### Fixing Large Effect QTL Increases Power for Detection of Additive Effects

The resolution and reproducibility of epic-QTL are further supported by additive QTL that do not map to known copper-resistance loci. Between the experiment with fixed chromosome I and the experiment with fixed chromosome VIII, we identified many overlapping selection-QTL peaks with locations of highest absolute z-scores within 1.5 kb of one another, indicating not only that there is high resolution and reproducibility but also that the underlying mechanisms of resistance do not strongly differ in the presence of one or the other fixed chromosome (Fig 3, S2 Table).

**Fig 3.**
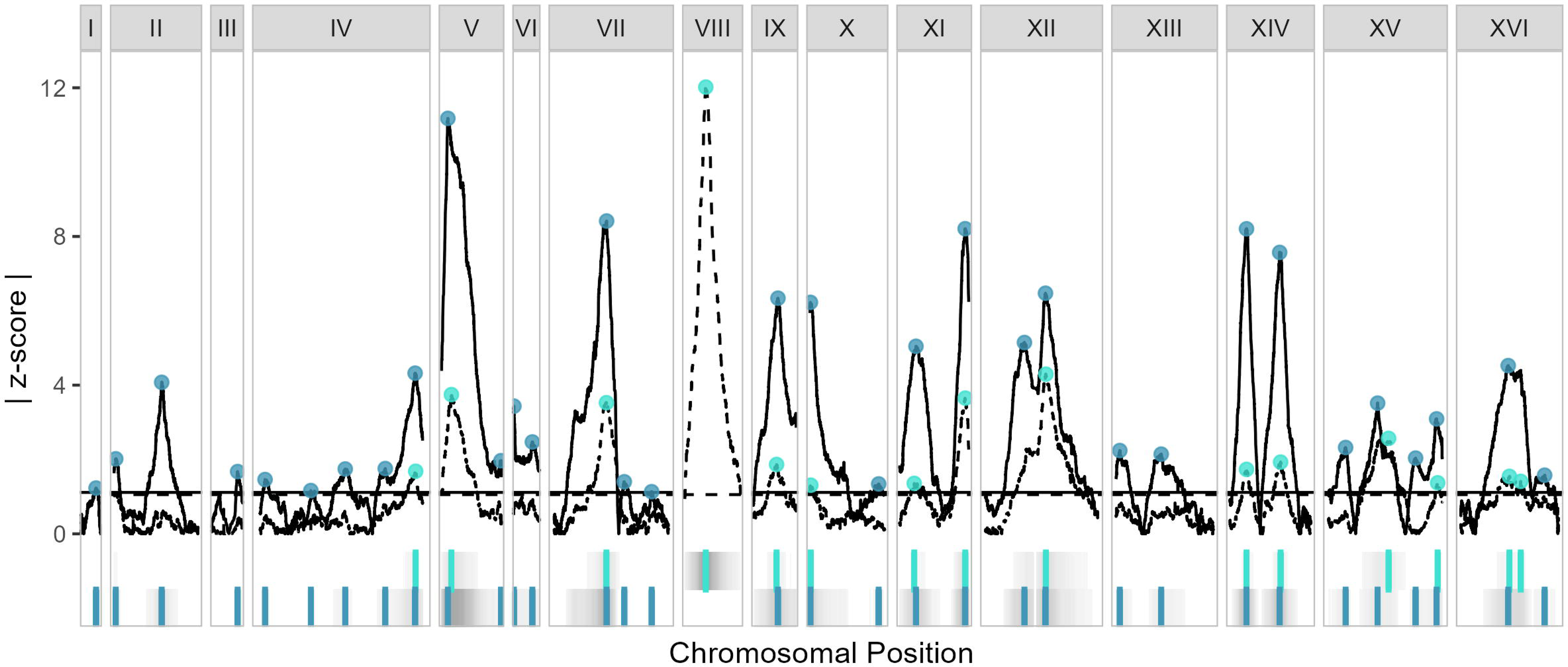
Consistent additive peaks identified in each fixed parental-chromosome experiment. Additive effects for the experiment with chromosome I fixed (dashed lines) and the experiment with chromosome VIII fixed (solid lines), shown as the absolute value of the selection coefficient’s z-score, with 5% FDR thresholds shown as horizontal lines of the same type. The only trace for chromosome I is solid, and the only trace for chromosome VIII is dashed, because of the fixed chromosome in each experiment. Peaks for each experiment are denoted as points on the traces as well as vertical lines below (fixed chromosome I as light blue, top strip, and fixed chromosome VIII as dark blue, bottom strip). In these strips, regions of significant z-scores are shown in gray, with intensities proportional to absolute z-score.

Having established that our method identifies additive QTL, we next investigated the result of chromosome fixation on additive effects. Removal of a major-effect QTL is expected to increase the power to detect QTL of lesser effect [41], so we investigated whether epic-QTL has an added benefit of increasing power when a major-effect QTL exists on a fixed chromosome. We found that indeed the majority of peaks in the fixed chromosome VIII experiment had z-scores of greater magnitude than the corresponding peaks in the fixed chromosome I experiment, and that several additional peaks were detected that were not called when chromosome I was fixed (Fig 3). On chromosome XII in particular, only a single peak is obvious in the fixed chromosome I experiment, whereas in the fixed chromosome VIII experiment, two peaks can clearly be differentiated (Fig 3). Taken together, our results for additive QTL demonstrate that epic-QTL has high power, resolution and reproducibility; that copper resistance in the Oak X Wine cross progeny shares at least some underlying mechanisms with other *S. cerevisiae* strains; and that removal of a major source of variance can increase power to detect other additive QTL.

### Magnitude epistasis is identified with chromosome VIII but not chromosome I

Reciprocal chromosome fixation effectively converts effects that are epistatic with that chromosome into additive effects that can be detected with high power by BSA. In the context of the epic-QTL logistic-regression model, these epistatic effects are identified as significant selection-by-background interactions.

We identify seven interactions with chromosome VIII: one each on chromosomes VII, X, XII, XIV, and XV, and two on chromosome XI (Fig 4, S3 Table). The z-score of each interaction peak is less extreme than that of its corresponding additive effect, but not always in the same direction as the additive effect, indicating that there is no consistent interaction effect (such as the Oak chromosome always amplifying Oak alleles’ selection effects). We did not identify any interactions at a 5% FDR with chromosome I (Fig 4A), despite observing >10 additive QTL. As chromosome I is the smallest of the *S cerevisiae* chromosomes, it is plausible that there would not be any interactions with the entire chromosome. This result provides evidence that epic-QTL does not generate spurious interactions, and that interactions are specific and not widespread across the genome.

**Fig 4.**
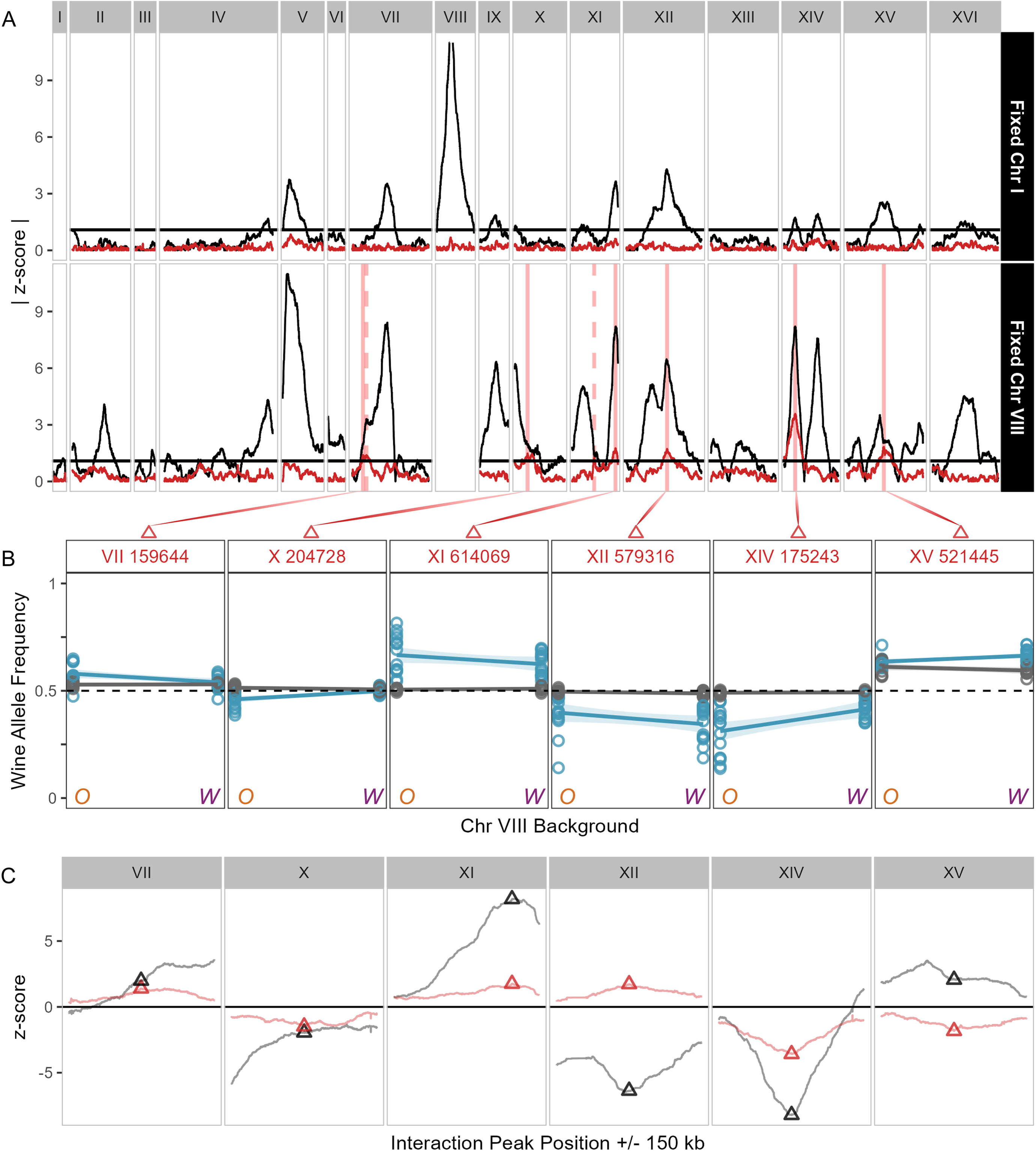
Interaction effects indicate magnitude epistasis. (A) Fixed chromosome I (top) and fixed chromosome VIII (bottom) absolute z-scores for CuSO_4_ experiments, with trace colors indicating additive (black) and interaction (red) effects.Vertical red lines indicate where an interaction peak has been called, with solid lines indicating the highest peak on a chromosome and dashed lines indicating a second significant peak. (B) Allele frequencies of the Wine allele at the maximum point of each interaction QTL per chromosome (indicated by solid lines in 4A and triangles in 4C), in each population. Oak (O) and Wine (W) fixed chromosome VIII backgrounds are on the horizontal axis, and the allele frequency of the Wine allele is on the vertical axis. Line crossing of selected (blue) and non-selected (gray) would indicate sign epistasis. (C) Comparisons of z-scores at interaction peaks (red triangles) with z-scores at corresponding additive peaks (black triangles).

The presence of an interaction can indicate different forms of epistasis. One possibility is sign epistasis, which in the case of epic-QTL would mean that changing the parent of the fixed chromosome changes the direction of the QTL effect at the interacting locus. An alternative possibility is magnitude epistasis, in which the QTL effect size at the interacting locus changes but the direction of the effect remains the same. In experimental designs such as standard BSA, where effects at any locus are averaged across a population, sign epistasis obscures phenotypic effect as the effects of allele pairs cancel each other out. Perfect sign epistasis, in which effect magnitude is equal but effect direction is opposite, is therefore impossible to detect by BSA without fixation of one allele in the pair to isolate the effect. In contrast, magnitude epistasis still produces an additive effect, although the effect size will be the average of that of each allele combination. To characterize our interaction QTL as exhibiting sign or magnitude epistasis with the fixed chromosome, we: 1) compared the allele frequencies of the highest points of the interaction peak (Fig 4C), and 2) scanned for loci above the 5% FDR threshold where the interaction z-score was more extreme than its corresponding additive z-score. Sign epistasis would require line-crossing of allele frequencies between parental backgrounds in the selected bulks, which we do not observe at interaction peaks (Fig 4B). We also do not find any significant interaction z-scores that are greater in magnitude than their selection effect (Fig 4C), which would indicate that the interaction obscures the additive effect through averaging. We therefore conclude that epistasis with chromosome VIII is magnitude and not sign epistasis.

Prior studies have identified interaction QTL for copper resistance in *S. cerevisiae* either by testing for interactions between additive QTL [4] or by barcoding combinations of alleles in natural variants [21,22]. We compared interactions identified with chromosome VIII in our study with those identified in these other studies, which utilized strains other than the Oak and Wine strains used here. We find that there is little overlap between identified loci (S2 Fig). There are, however, consistent additive effects at specific loci such as *CUP1*, and indeed most interactions with chromosome VIII from the other studies did involve the *CUP1* locus. These results suggest that interactions are between specific alleles not necessarily present in all crosses, which could be corroborated with additional epic-QTL experiments with different parental strains.

Further evidence that interaction effects are idiosyncratic is that not every additive locus interacts with the fixed chromosome, and there is no correlation in the direction or magnitude of effect of the interaction compared to additive effect (Fig 4C). The majority of interaction peaks (4 of 7) were within 6 kb of the additive peak maxima, indicating that the interactions with chromosome VIII are likely with the same loci that produce additive effects. Of the remaining interaction peaks, two (the one on chromosome VII and the lesser one on XI) overlap with what appear as shoulders of additive peaks, suggesting that multiple QTL occur within that peak, and that the second largest is interacting with chromosome VIII. The peak on chromosome XV might also correspond more with the shoulder of the additive peak. The interaction peak and corresponding additive peak shoulder on chromosome VII are close to the *CUP2* gene, which induces transcription of *CUP1* [35]. *CUP2* could thus be a candidate for allele substitution to confirm the interaction with *CUP1* on chromosome VIII. Overall, the co-occurrences of interaction peaks with additive peaks indicate that interactions occur primarily with loci that have an additive effect, and the lack of interactions at every additive peak, or even the most significant additive peaks, suggests that interactions are specific and idiosyncratic.

## Discussion

Identifying epistatic contributions to trait variation presents challenges caused by the combinatorial nature of interactions and their potential to obscure effects of individual alleles. Methods to detect pairwise interaction effects suffer from low statistical power due to the number of pairwise comparisons across the genome to test. High power can be achieved by BSA experiments, but averaging over combinations of alleles precludes analysis of interactions. Because of these challenges, approaches to predicting the effect of genotype often minimize the contribution of epistasis to phenotypic variance, by first (or only) fitting additive effects [9]. However, although the effects of epistasis might be statistically underestimated, specific examples of epistasis have been observed across species and traits such as in tomato size and fruit number [42,43], corn kernel coloration [44], and mouse coat color (39, reviewed by 12). In non-natural variants such as deletion mutations or engineered hypomorphs, interactions have also been quantified; using high-throughput approaches, Costanzo *et.al.* found ∼900,000 interactions among 23 million double mutants representing nearly every pair of genes in the yeast genome [24], and Hale *et al.* found that over one quarter of 1721 CRISPR-interference perturbations showed growth effects that depend on genetic background [46]. We developed epic-QTL with the aim of bringing the power of a high-throughput, DNA sequencing-based approach to the identification of interactions between natural genetic variants. Specifically, we circumvented the statistical problem of pairwise testing of natural variants by reciprocally fixing a single chromosome from each of two genetic backgrounds in a cross between divergent yeast strains. With the high power afforded by BSA, we observe magnitude epistasis with additive QTL but no evidence of cryptic sign epistasis.

We first verified our method by comparing additive QTL identified by epic-QTL, both between experiments with chromosome I or chromosome VIII fixed and between our study and those conducted on copper resistance in other strains. Known drivers of copper resistance, *CUP1* and *FRE1*, were identified as additive QTL in all experiments where the loci were allowed to segregate. Additive peaks colocalized between the experiment with chromosome I fixed and the experiment with chromosome VIII fixed, but on average those identified in the experiment with fixed chromosome VIII were stronger, very likely due to the fixation of a major QTL. In comparison with other studies of copper sulfate resistance, several additive QTL were found in common, but many were not, suggesting strain-specific differences in which alleles are present that impact resistance.

The key strength of epic-QTL is identification of interactions with fixed chromosomes at high statistical power. We next compared interactions found by epic-QTL to those identified for copper resistance using different strains, different methods and smaller sample sizes [2,19,21,22], finding that although there is some overlap in regions containing additive QTL, the majority of interactions are not common between studies. This finding speaks to the importance of more high-powered studies of interactions and potentially also suggests that interactions are idiosyncratic in that they are dependent on the specific alleles found in divergent strains. Because we detected more additive than interaction QTL, not every detected additive QTL in our study contained interaction QTL. Moreover, interactions did not necessarily cluster only under the highest additive peaks. This finding further supports that the interactions are idiosyncratic, rather than globally caused by nonlinear transformation of the value of a latent trait with additive genetic architecture [14]. One key overlap with another study to note, however, is *CUP1* as a hub for interactions on chromosome VIII [22]. That interactions occur with chromosome VIII but not chromosome I, and that interactions occur primarily with additive QTL, suggests that our interactions might be with *CUP1*. The logical next step would be to fix the chromosomes found to interact with fixed chromosome VIII, allowing chromosome VIII to segregate, and thereby to identify the interaction partner(s) on this chromosome.

The interaction QTL identified here provide evidence of three compelling trends that imply a middle ground between completely idiosyncratic epistasis and highly predictable global epistasis: 1) interaction peaks commonly colocalized with additive peaks, 2) magnitude epistasis was observed whereas sign epistasis was notably absent, and 3) interaction peaks differ between studies and are not driven by strength of an additive peak. Sign epistasis has the potential to completely obscure additive effects, and thus the lack of sign epistasis implies that interactions occur primarily with loci that already have an effect. Precise colocalization of additive and interaction maxima suggest that the additive QTL interacts with the fixed chromosome. These trends corroborate the conclusion by Bloom *et al.* [4] that interactions occur primarily between additive QTL, and they validate the approach of reducing the search space by only looking at interactions between additive QTL as done by Nguyen Ba *et al.* [21].

Sign but not magnitude epistasis can influence available evolutionary trajectories by creating valleys between peaks on a fitness landscape [11,13]. Examples of specific reciprocal sign epistasis, for example mutually exclusive adaptive mutations in the *HXT6/7* and *MTH1* genes during experimental evolution under glucose limitation, have been shown to cause fitness valleys [47]. Sign epistasis involving mutations in specific interaction hubs has also been observed [25], but the general prevalence of sign epistasis is yet to be determined. epic-QTL offers an unbiased, powerful method for detecting sign epistasis. The lack of sign epistasis detected here therefore indicates that it is relatively rare, at least for the trait, fixed chromosomes, and parental strains we studied.

It will be important to see whether the trends we observed generalize to other fixed chromosomes and other complex traits. A complete set of fixed-chromosome strains would yield an additional benefit of very rapid assessment of the extent to which epistasis impacts variation in a trait, in that a completely additive genetic architecture would mean that the sum of chromosome effects should equal the difference between the parental strains. Such a test only requires chromosome-substituted strains, not segregants from them. When this kind of test was applied in mice and rats to a large number of anatomical and physiological traits, it revealed much evidence for epistasis [48].

A widespread lack of sign epistasis would greatly aid in identifying interactions in studies where experimental manipulations of genotypes are not possible. Genome-wide association studies in particular face challenges with respect to the number of subjects relative to the number of possible interactions between markers, and so a variety of methods have been proposed to avoid exhaustively searching all possible pairs of polymorphisms [49]. Prioritizing interactions between additive loci rather than every possible combination could therefore be a principled approach to increase the power to detect interaction effects. Characterizing interaction effects in humans should in turn allow for greater precision in predicting response to medication and susceptibility to disease, as well as facilitate further understanding of the molecular mechanisms of complex traits. In all, our characterization of epistasis not only enhances our understanding of genetic interactions in model organisms but also has implications for improvement of applications in personalized medicine and trait prediction across diverse species.

## Materials and Methods

### Conditional Chromosome Strain Construction

Two divergent haploid yeast strains, one oak tree-derived (NCYC3631, MATα) and one wine barrel-derived (NCYC3591, MATa) [29], were each engineered for chromosome removal from diploids and later selection of haploid progeny. To facilitate crosses, strains were modified by homologous recombination at *URA3*, *HO*, and *MAT*α using standard lithium-acetate-based transformation. *URA3* prototrophy was restored (and *KanMX* removed) in wine-derived strains by insertion of *URA3* sequence amplified from the wine-strain parent (BC187). The positive-selection marker *hph*, which confers resistance to hygromycin, was removed from the *HO* locus of each parental strain to allow the marker’s use elsewhere. To remove *hph*, we conducted Delitto Perfetto excision [50] using a cassette that allows positive selection for nourseothricin resistance (*NatR*) and negative selection against HSV-*TK*, which encodes the thymidine kinase from herpes simplex virus and thereby confers sensitivity to the pyrimidine analogue 5-fluorodeoxyuridine (FuDR) [51]. Importantly, two identical sequences (*cyc* terminators) flank this cassette and thereby lead to spontaneous excision of the cassette by homology-based repair [50]. Transformants were selected for presence of *NatR* and checked for loss of *hph*. We then counter-selected using FuDR to select for removal of the cassette and resulting loss of both *NatR* and HSV-*TK*.

To select for haploids of one mating type in bulks to be sequenced, we engineered the *MATα* locus to contain the HSV-*TK* negative-selection marker for removal of any cell that is not a mating-type a haploid. *MATα* contains a bidirectional promoter driving the expression of the *α1* and *α2* genes, both of which must be active for the cell to sporulate [52]. The negative-selection marker was inserted upstream of *α1* and its promoter, and the *STE3* promoter (*pSTE3*), which is activated by the transcriptional co-activator encoded by *α1*, was placed upstream of *α2* [53]. Thus, the selection cassette was placed in a way that maintains expression of both *α1* and *α2* in mating-type α haploids and in diploids. As a result, the cells retain the ability to mate with mating-type a cells as haploids and to sporulate as diploids, but both diploids and haploid mating-type α haploids are sensitive to FuDR. For this transformation we used the positive-selection marker *hph*. We transformed a cassette containing MATα homology and the positive- and negative-selection markers into the Oak α strain, and then replaced *hph* with *NatR* using lithium acetate homology-directed transformation, allowing use of *hph* in later transformations.

For chromosome elimination by driving transcription through the centromere, we constructed an inducible-promoter cassette by using overlapping PCR to combine the HSV-*TK* and *hph* markers from Alexander *et al.* [51], the *GAL1* promoter (*pGAL1*), and regions homologous to the yeast genome adjacent to the centromere. The centromere-region primers matched the primers used by Reid *et al.* [30] or matched sequences very close nearby. These cassettes were transformed into either chromosome I or chromosome VIII in either the Oak α or Wine *a* strains. All strains are detailed in supplemental data (S4 Table). These haploid strains are referred to as the background (Wine a or Oak α) plus [I] or [VIII] to indicate the chromosome that can be eliminated.

Wine a[I] and Oak α were mated to produce diploids, and Wine a and Oak α[I] were mated to produce diploids capable of producing the reciprocal chromosome fixation (Fig 1A). Diploids were selected by ability to grow in both the absence of uracil and the presence of nourseothricin, and ploidy was confirmed by flow cytometry using DNA staining with propidium iodide on the Cytek Aurora. To fix chromosome I, diploids were then plated onto medium containing 2% D-Galactose (and lacking glucose) to induce the *GAL1* promoter’s transcription through the centromere of the conditional chromosome. Although we did not use markers to select for the loss of a chromosome, plating on galactose and restreaking from single colonies was effective for yielding chromosome loss. Colonies from the restreaked plate were checked by Sanger sequencing to identify a loss of the conditional chromosome (i.e. by loss of heterozygosity on that chromosome), then grown in liquid media and frozen as clonal aneuploid (2N–1) strains, which we term Oak I or Wine I to indicate the parent of origin of the remaining chromosome I in these otherwise hybrid strains. Populations of ∼10^7^ clonal aneuploids were sporulated using 2% potassium acetate for ∼10 days at room temperature (25 °C) to produce approximately 4X10^7^ segregants with identical chromosome I of either Oak or Wine background but segregating parental genotypes at all other loci. To select for haploids above and beyond the MAT*α* selection, spores were vortexed for 10 minutes in 50% diethyl ether to kill unsporulated diploids [54]. Surviving spores were rinsed twice in sterile water, then germinated by incubation in 1 mg/mL Zymolyase (Z100T) for 12 minutes before culturing in each experiment. The process was completed identically for the reciprocal fixations of chromosome VIII.

### Concentration Determination for Selection

We tested approximate concentrations of copper for selection by growing cells in media of different concentrations for 24 hours (matching the experiment time) and then counting the cells either by automatic cell counter (ThermoFisher Countess™ II) or by manual counting on a hemocytometer. From there, we chose smaller ranges of doses that would provide between 1% and 10% survival. After conducting experiments, cell counts in some experiments were checked by hemocytometer to confirm that experimental treatments were achieving the intended amount of selection.

### Bulk Segregant Analysis

Pools of segregants with reciprocal fixed chromosomes (Oak I and Wine I, or Oak VIII and Wine VIII) were grown in parallel for 24 hours in either YPD or YPD + copper sulfate at doses determined in each chromosome-fixed bulk population to produce between 1% and 10% survival. The control cultures grown in YPD were seeded at 10% of the starting density for the selected group. Each of the four bulk categories was grown in duplicate. After 24 hours, each of these bulks was harvested by spinning down cells separately and extracting DNA either by phenol-chloroform-isoamyl alcohol extraction and ethanol precipitation [55], or by Qiagen DNeasy Blood and Tissue Extraction Kit 69506.

### Sequencing

After DNA was harvested, single-end libraries for DNA sequencing were prepared according to Baym *et al.*’s protocol [56] using Illumina kit FC-131-2001 and KAPA HiFi amplification kit KK2612. Fragments were selected to be 200–600 bp by AMPure XP bead cleanup (Fisher NC9959336), and pooled based on concentrations determined by LifeTechnologies Qubit 3.0 and Roche 480 LightCycler qPCR. Libraries were then sequenced on an Illumina NovaSeq 6000, with an average per-bulk depth of ∼70X. Reads were aligned to the S288C reference genome R64 using BWA-MEM [57], sorted using PicardTools [58], and then underwent quality control using Genome Analysis Toolkit [59] command *BQSR*. Variants were called using GATK *HaplotypeCaller* [60] as described by Mansfield and Grumet [61].

### epic-QTL Analysis

VCFs were converted to tables of reads to analyze in R using GATK’s *VariantsToTable* function. We first removed all loci with >2 alternate alleles to control for mapping quality issues, then assigned each alternate allele to the corresponding parent based on SNP location from whole genome sequencing of the parent strains [62]. To increase our signal-to-noise ratio and take advantage of the inherent correlation of nearby genomic positions in a single F2 cross, we smoothed the read counts in rolling windows of 200 SNPs using a Gaussian weighted mean.

To partition the effects of fixed-chromosome parent of origin, selection, and their interaction, we performed logistic regression on these counts for each smoothed position using the following formula:

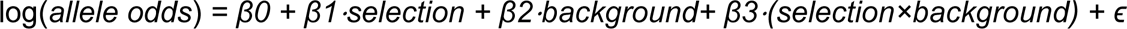

where *selection* is the presence or absence of copper sulfate in the culture media, *background* is the parent of origin of the fixed chromosome, and the *selection×background* term captures the dependence on background of the selection effect, i.e. epistasis. We then used the z-scores of each effect to identify additive (*selection*) and epistatic (*selection×background*) QTL. Absent any effects of the fixed chromosome’s parent of origin on segregation of the other chromosomes or on growth in the absence of copper sulfate, we do not expect to see a significant background effect. For this reason, we set the contrasts of the selection variable to 0 (copper absent) and 1 (copper present), and for the background variable to –0.5 (Wine) and 0.5 (Oak). Outputs of this analysis are available in supplemental data (S1 Table).

To establish our statistical threshold for interactions, we conducted permutations of allele frequency ratios by shuffling positions within each parent and replicate in the non-selected bulks. This permutation approach is based on the assumption that, in the absence of selection, there would be no loci associated with the trait. For the permutations, we excluded chromosomes I and VIII from all experiments to remove the effect of the fixed chromosomes, and removed chromosomes III and V, which contained engineered regions not identical in the two parents. We then assigned bulks randomly to these permuted values according to the structure of the original dataset (the same number of replicates in each group) and calculated our selection, background, and interaction effects for each. These effects drawn from the bulks of two different parents form the null distribution of z-scores of interaction effects for each experiment. All analyses employ a false discovery rate threshold of 5%.

Peaks of absolute z-score (QTL locations) were called by identifying the maximum points within chromosome segments bounded by chromosome ends or local minima, defined as positions where the local slope of the absolute z-score trace shifted from negative to positive. The local slope was calculated within sliding windows of 700 SNPs by linear regression within the window. The highest point in that segment, if above the 5% FDR, was then called as the peak. Additive and interaction peak locations are available in supplemental data (S2 and S3 Tables).

## Acknowledgements

This work was supported by National Institutes of Health grant R35GM148344 (to MLS), the NYU Biology Summer Undergraduate Research Program (JKV), and the NYU Graduate School of Arts and Science Dean's Dissertation Fellowship (CB). This work was supported in part through the NYU IT High Performance Computing resources, services, and staff expertise. We acknowledge the Zegar Family Foundation for their generous support. We thank the NYU Center for Genomics and System Biology Genomics Core for their assistance and resources. We thank members and alumni of the Siegal lab for input on the paper and discussion about the project.

## Supporting information captions

**S1 Fig. Survival percentage by strain in CuSO4.** Boxplots of survival percentages of Oak (NCYC3631) and Wine (NCYC3591) haploid strains at different concentrations of copper sulfate are shown. Replicate experiments carried out on different dates are represented by different shapes.

**S2 Fig. Comparison with copper sulfate interaction peaks from previous studies.** Interactions with chromosome VIII identified in our study (shown in black trace of the absolute z-score by genome position) are compared with interaction QTL involving chromosome VIII identified in other copper-resistance studies, which used both different experimental designs and different strains. Beneath the trace from this study, NB (magenta) indicates interaction QTL as identified by Nguyen Ba et al [21], M (turquoise) represents interaction hotspots between strains from Matsui et al. [22], and B (dark blue) represents interaction QTL as identified by Bloom et al [4]. Red vertical lines indicate significant peaks called in our study. *CUP1* is indicated by the vertical light blue dotted line on Chromosome VIII.

**S1 Table. Logistic regression output**. Direct output from logistic regression on sequencing data for each divergent SNP that passes quality filters (see github for code). Each row (numbered in column A) records the effect term in the model (Background, Bulk, Interaction, or intercept (column B, “label”); the chromosome fixed in the experiment (column C, “chr”); the chromosome containing the SNP (column D, “CHROM”); the chromosomal position of the SNP (column E, “POS”); the z-score of that SNP in that experiment and for the specified effect (column F, “zscore”); and the threshold value for a 5% FDR for the specified effect and experiment (column G, “q5”).

**S2 Table. Additive-effect peaks**. Peaks called for the Bulk effect. Each row records the chromosome fixed in the experiment (column A, “CSS”), the location of the peak SNP as its chromosome (column B, “CHROM”) and chromosomal position (column C, “POS”), and the z-score of the peak SNP (column D, “zscore”).

**S3 Table. Interaction peaks**. Peaks called for the Interaction effect. Columns are as in S2 Table.

**S4 Table. Yeast strains constructed and used in experiments**. Backgrounds, genotypes, and aliases of all strains used, including those representing intermediate steps to construct strains for epic-QTL experiments. MLS.CB refers to strains made in this study; NCYC refers to strains from the NCYC collection [29].

