## Supplementary figures and images for "Epistasis and cryptic QTL identified using modified bulk segregant analysis of copper resistance in budding yeast"

### S1 Fig

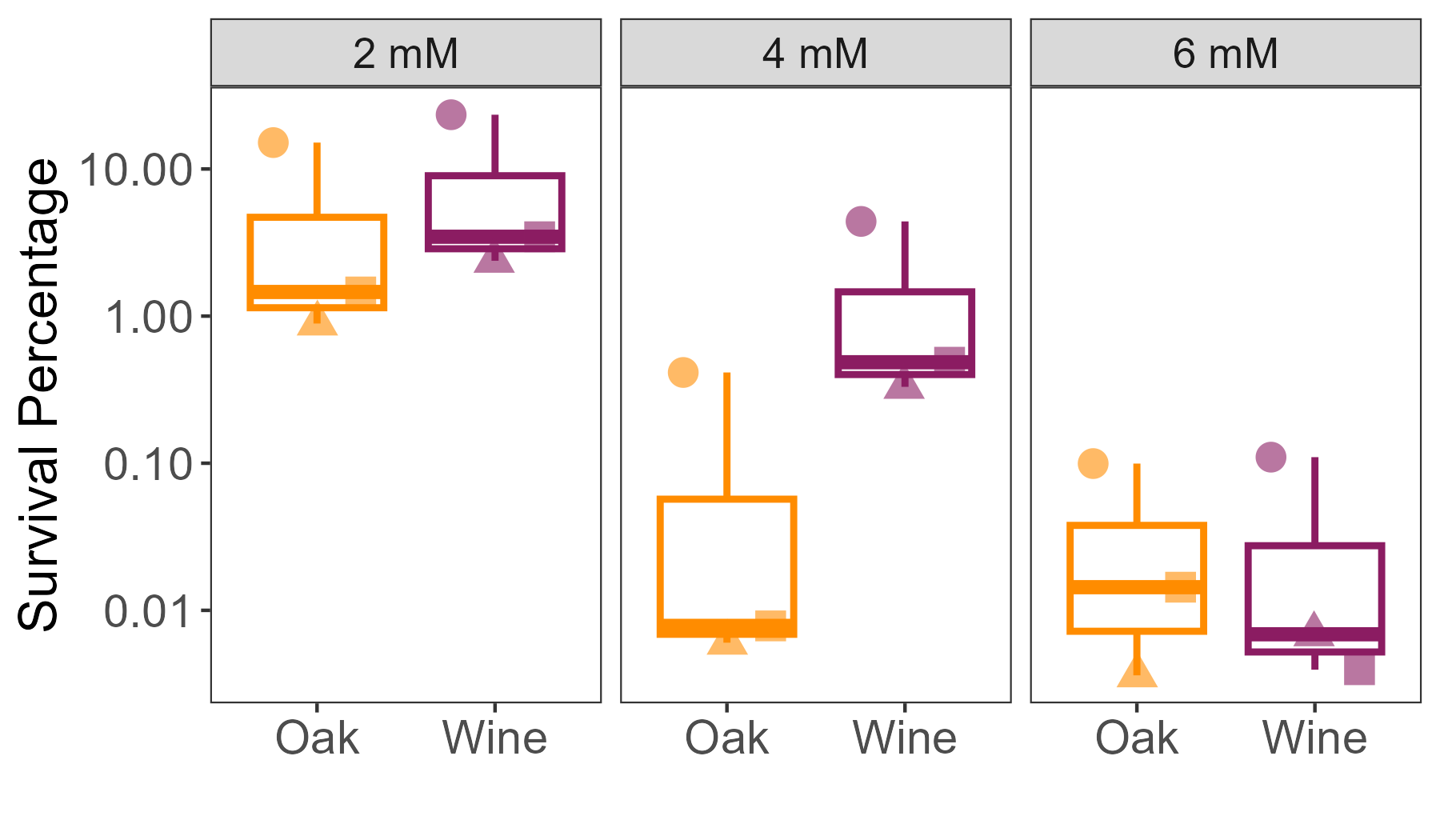

### S2 Fig

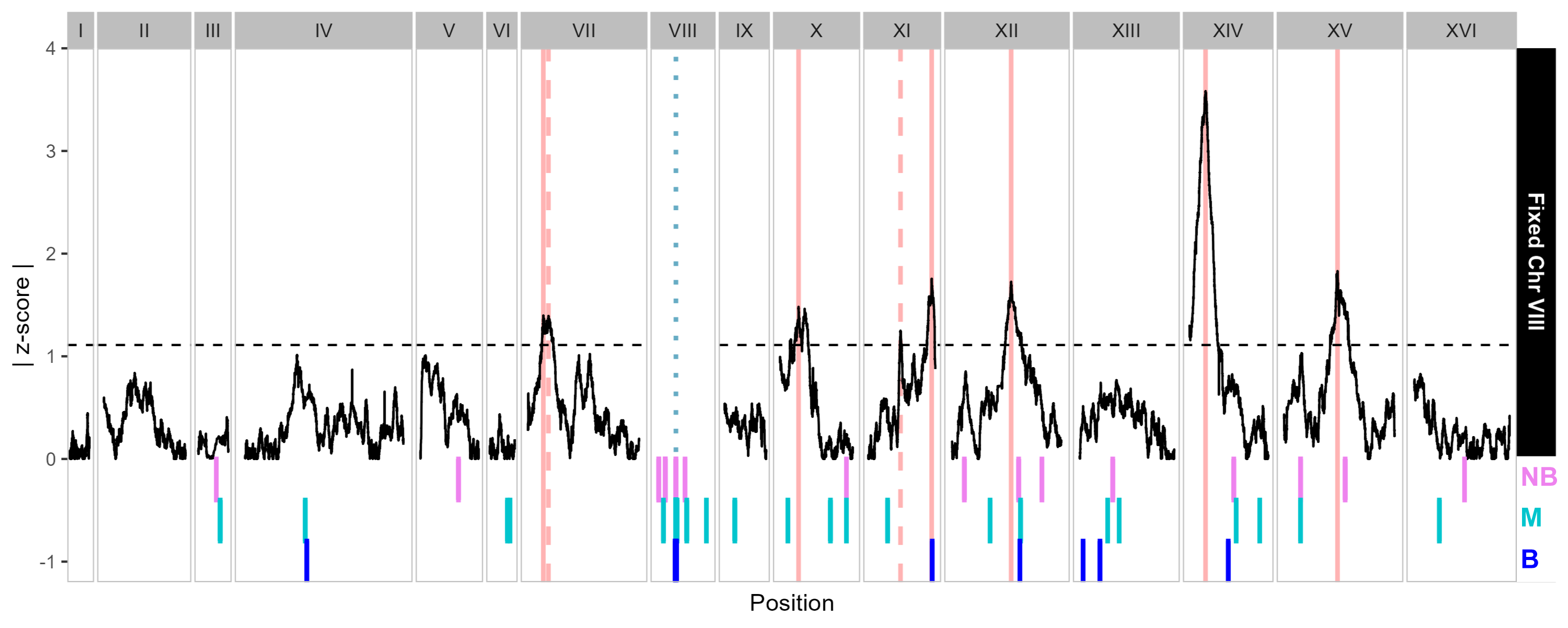
